# Coordinated horizontal transfer of multiple genes assembles a carotenoid biosynthesis pathway in aphids

**DOI:** 10.1101/2025.03.24.644864

**Authors:** Rong Hu, Jun Wu, Siying Li, Peiyu Yang, Gang Wu, Changying Niu, Shuai Zhan, Yazhou Chen

**Author notes:** Correspondence: Yazhou Chen.

## Abstract

Horizontal gene transfer (HGT) plays a crucial role in genome evolution, especially when it enables the acquisition and assembly of multi-step biosynthetic pathways. Here, we investigate the evolutionary origins of carotenoid biosynthesis genes in aphids to determine whether multiple functionally related genes were acquired through HGT. We analyzed carotenoid biosynthesis genes in 23 aphid genomes based on homologs in plants, fungi, and bacteria. Phylogenetic analyses revealed that *Geranylgeranyl pyrophosphate synthase* (*GPS*), *Phytoene synthase* (*PS*), and *Carotenoid desaturase* (*CD*) were acquired via HGT from fungi by ancestral insect species, while *Carotenoid cleavage oxygenase* (*CCO*) appears to be a native insect gene. Most insect genomes contain two *GPS* copies, likely resulting from independent HGT events, whereas aphid genomes exhibit extensive duplication of *PS* and *CD*, a pattern uncommon in other insects. Expression analyses across aphid species with distinct pigmentation showed that these genes are broadly transcribed with substantial variability in expression levels. In *Myzus persicae*, comparative expression analysis between reddish and greenish clones, as well as a green-reddish clone with green and red color polymorphism, revealed that *PS-4390* is a novel contributor to red pigmentation in *M. persicae*, in addition to *CD-4400*, a homolog of the *tor* gene in *Acyrthosiphon pisum*. These findings provide strong evidence that HGT can introduce multiple functionally related genes into recipient genomes, allowing them to be co-opted into a functional biosynthetic pathway.

## Introduction

Horizontal gene transfer (HGT), the acquisition and integration of foreign DNA fragments into the recipient genomes, is a fundamental driver of adaptation and evolution (Treangen and Rocha 2011; Schönknecht et al. 2014; Blakely 2024). Initially, HGT was thought to be predominantly restricted to prokaryotes, but it has now been widely observed across both prokaryotic and eukaryotic lineages, contributing to genomic innovation and ecological adaptation (Koonin 2016; Keeling 2024).

In bacteria, HGT facilitates rapid adaptation to environmental challenges by enabling the uptake of advantageous genes from external sources (Johnston et al. 2014; Power et al. 2021). For instance, approximately 12% of the core *Bacillus subtilis* genome undergoes gene transfer within just 200 generations, significantly enhancing their population fitness (Power et al. 2021). Many HGT-acquired genes in bacteria encode catabolic enzymes, expanding the metabolic repertoire of recipient species for nutrient utilization (Goyal 2022). HGT in eukaryotes is more complex but has been increasingly documented (Van Etten and Bhattacharya 2020). Large-scale HGT events have been identified in fungi such as *Armillaria* species. These horizontally transferred genes enhance their pathogenicity and lignin degradation in host plants (Sahu et al. 2023). Similarly, in the parasitic *Cuscuta* plants, approximately 108 HGT events have been detected, enhancing their resistance and small RNA production to interact with host plants (Yang et al. 2019).

In insects, HGT is more frequent and complicated than previously thought (Li et al. 2022). Across 218 insect genomes, more than 1400 HGT-derived genes have been identified (Li et al. 2022). Insects acquired genes originated from diverse sources, including bacteria, viruses, fungi, plants, and even other animals (Liu et al. 2023; Xing et al. 2023; Yadav et al. 2024). Microbial-derived toxin genes acquired via HGT have been found in numerous insect species, bolstering immune defense against natural enemies (Verster et al. 2021). Additionally, extensive gene transfer has been observed between *Chordodes* nematomorphs and their mantis hosts, where over a thousand genes appear to have been exchanged, potentially facilitating nematomorphs to manipulate hosts (Mishina et al. 2023).

Intriguingly, insects have also acquired genes directly from the host plants, likely through prolonged and intimate interaction. The whitefly *Bemisia tabaci* has integrated multiple plant-derived genes that contribute to novel physiological functions, enhancing host colonization (Xia et al. 2021). Among these, genes encoding malonyltransferases and Δ12 desaturase have been acquired from plants, conferring potential metabolic advantages (Xia et al. 2021; Gong et al. 2024). Similarly, a thaumatin-like protein gene has been horizontally transferred from plants into both *B. tabaci* and the closely related species *Trialeurodes vaporariorum*, suggesting a broader role for plant-to-insect HGT in insect-plant interactions (Hu et al. 2025).

Although numerous HGT-derived genes have been identified across diverse species, the function of most remains unclear. In many cases, horizontally transferred genes act as modifiers of existing biological processes. However, some HGT-acquired genes are involved in entirely novel and complex biological processes that are absent in related species (Danchin et al. 2016; Tapia et al. 2023; Yadav et al. 2024). This suggests that multiple genes required for such complex processes may have been acquired through HGT. For instance, in *Bemisia tabaci*, two sequentially acquired HGT-derived genes, *BtUCA* and *BtAtzF*, coordinately participate in nitrogen metabolism by facilitating the conversion of urea into amino acids, extending the existing nitrogen metabolic pathway (Zhang et al. 2024). A few genes, such as *MurF* and *DapF*, acquired by *Planococcus citri* via HGT from bacteria, have been shown to function in coordination with the insect’s own genes and those of its symbionts to form a mosaic Peptidoglycan (PG) biochemical pathway(Bublitz et al. 2019).A striking example of HGT involving multiple genes is the carotenoid biosynthesis pathway—which is typically absent in animals but widespread in plants, fungi, and bacteria (Heath et al. 2013; Clark and Lampert 2018; Ding et al. 2022). In carotenoid-producing organisms, the pathway begins with the condensation of geranylgeranyl pyrophosphates (*GGPP*) to phytoene, catalyzed by phytoene synthase (Rohdich et al. 2001; Nisar et al. 2015; Avalos et al. 2017). Phytoene is then converted into intermediates, such as ζ-carotene and lycopene, through sequential desaturation steps. These early steps are conserved across plants, fungi, and bacteria and involve key enzymes, including geranylgeranyl pyrophosphate synthase (*GPS*), phytoene synthase (*PS*), and carotenoid desaturase (*CD*) (Rosas-Saavedra and Stange 2016; Kovács et al. 2003; Giraud et al. 2004; Avalos et al. 2017). The latter stages of carotenoid biosynthesis diverge among species, resulting in lineage-specific variations in carotenoid composition.

Aphids, however, possess the ability to synthesize carotenoids, a trait attributed to the horizontal acquisition of multiple fungal genes, including those encoding carotenoid synthase/cyclase and carotenoid desaturase (Moran and Jarvik 2010; Nováková and Moran 2012, Cobbs et al. 2013, Trissi et al. 2023, Ge et al. 2024). Given that carotenoid biosynthesis is a multi-step process requiring several enzymes (Rosas-Saavedra and Stange 2016), it is plausible that additional carotenogenic genes in aphids may also have been acquired through HGT. Moreover, these acquired genes appear to have undergone extensive duplications, with some duplicated copies functionally implicated in carotenoid biosynthesis (Moran and Jarvik 2010, Grbić et al. 2011, Bryon et al. 2017, Takemura et al. 2021, Trissi et al. 2023, Ge et al. 2024). For example, the carotenoid desaturase gene *tor*, which is a key determinant for red/green pigmentation in the *Acyrthosiphon pisum*, exists in two copies in *Myzus persicae* (Moran and Jarvik 2010). However, only one of these copies appears to be involved in red coloration in *M. persicae* (Trissi et al. 2023; Ge et al. 2024). With many carotenogenic genes actively expressed in aphids (Trissi et al. 2023), these raise a central evolutionary question: how have horizontally acquired and duplicated carotenoid biosynthesis genes been functionally integrated, diversified, and retained in aphid genomes.

In this study, we investigate carotenogenic genes in aphids by examining the homologs of plants, fungi, and bacteria. Our analysis revealed that genes in core carotenoid biosynthetic pathways (*GPS*, *PS*, *CD*, and *CCO*) are present in most aphid species and some other insects. Phylogenetic analysis indicates that *GPS*, *PS*, and *CD* were acquired from fungi via horizontal gene transfer (HGT), whereas *CCO* is of native insect origin. These HGT-acquired genes have undergone extensive duplication during aphid diversification and exhibit variable expression patterns across aphid species with distinct body colors. By comparing gene expression in differently pigmented *Myzus persicae*, we identified *PS-4390* as a previously unrecognized component of aphid carotenoid biosynthesis. This study systematically elucidated the carotenoid biosynthesis pathway in aphids and introduced a novel regulatory gene to aphid body color.

## Results

### Carotenogenic genes widely exist in insect genomes

Carotenoid biosynthesis pathways are present in plants, bacteria, and fungi. In total, 26 genes have been reported to participate in the pathways, 11 from plants, 7 genes from bacteria, and 8 from fungi (Supplementary fig. S1, Supplementary table S1). The genes in the upstream pathways, including *Geranylgeranyl pyrophosphate synthase* (*GPS*), *Phytoene synthase* (*PS*), and *Carotenoid desaturase* (*CD*) (Fig. 1A), are conserved among different kingdoms. Genes in the downstream pathways are more kingdom-specific (Fig. 1A, Supplementary fig. S1).

**Fig. 1.**
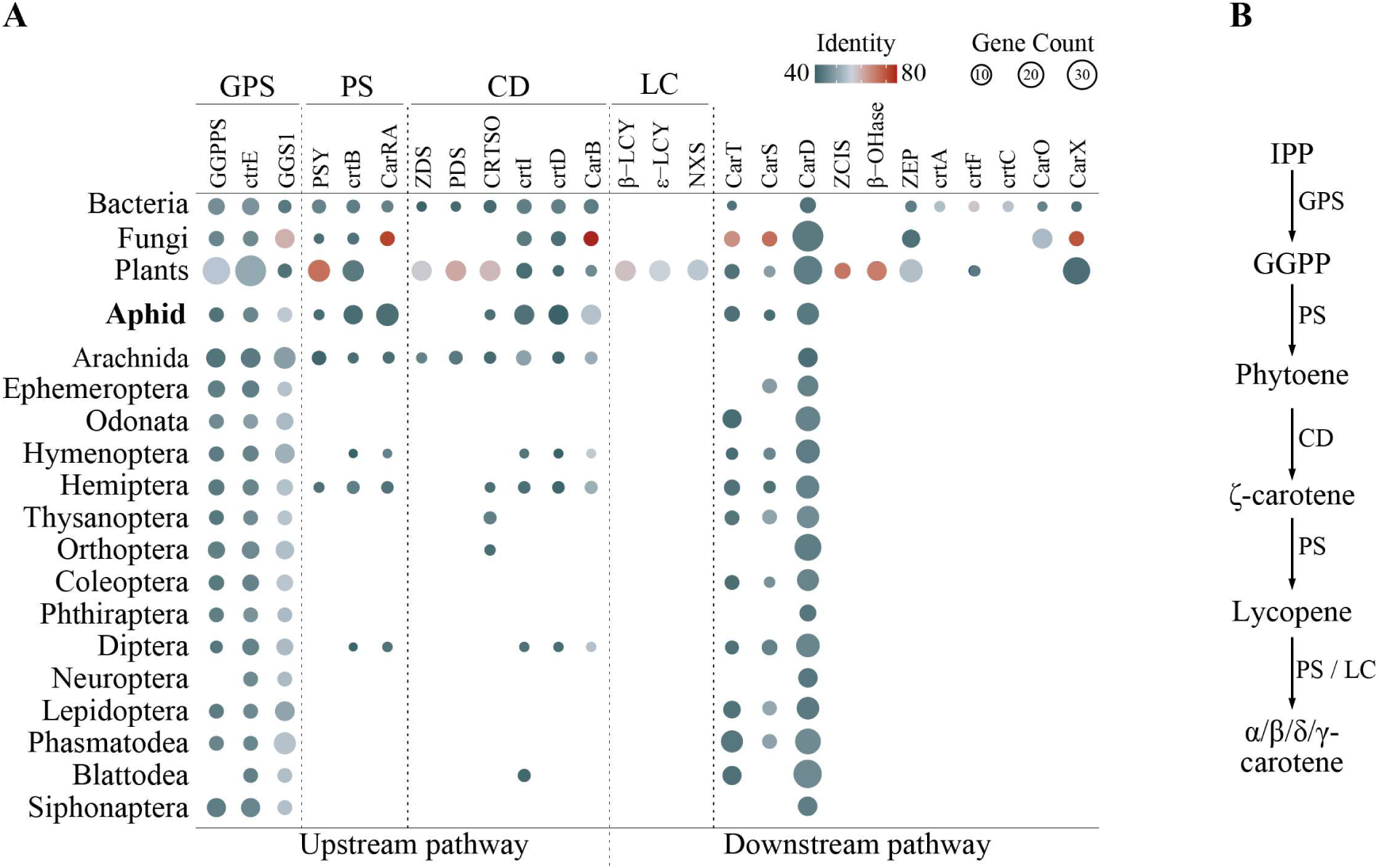
Carotenogenic genes are widespread in aphid and other insect genomes. (A) Distribution of carotenogenic genes across the plant, fungi, bacteria kingdoms, as well as the insect class. Bubble size represents the average gene count per species (total gene count/total species). Gradient color indicates sequences with >40% BLASTp identity. Genes belonging to the same gene family and sharing an identical Pfam domain are grouped together. Gene abbreviations (*GPS*, *PS*, *CD*, *LC*) correspond to the uppercase initials of their full names. The upstream pathway represents a conserved pathway across the three biological kingdoms, while the downstream pathway is species-specific. For pathway details, see Supplementary fig. S1. (B) Putative upstream pathway for insect carotenoid biosynthesis. The pathway in insects was inferred based on references from the three kingdoms.

To investigate the carotenoid biosynthesis pathway in aphids, we performed a reciprocal BLASTp in which the 26 genes were searched against reference protein sequences retrieved from 23 aphid genomes. The homologs identified were validated by the presence of conserved Pfam domains, resulting in the identification of five carotenogenic genes in different aphid species (Fig. 1A, Supplementary table S2). These included three upstream genes (*GPS*, *PS*, and *CD*) and two downstream genes in the fungal pathway (*carT* and *carD*) (Fig. 1). These findings suggest the entire carotenoid biosynthesis pathway likely exists in the aphid genomes (Fig. 1B).

To determine whether the pathway also exists in other insects, we extended the BLASTp analysis to 265 additional insect species. The homologs of *GPS*, *PS*, *CD*, *carT*, and *carD* were identified in most insect species (Fig. 1A, Supplementary table S2). The homologs of *carT* and *carD* in insects were annotated as *Carotenoid isomerooxygenase* (also known as *Carotenoid cleavage oxygenase, CCO*) (Liu et al. 2024b), and *Aldehyde dehydrogenase* (*ALDH*) in insects, respectively. *ALDH* genes were excluded from the rest of the analysis as they are unlikely to be relevant to the carotenoid biosynthesis pathway. These findings indicate that the carotenogenic genes are widely distributed among insects, although some components may be less conserved.

### Carotenogenic genes were acquired from fungi by the ancestor species and duplicated in aphids

Aphid *tor* genes (*CD*) are known to have been acquired through HGT from fungi (Moran and Jarvik 2010). To investigate whether other carotenogenic genes identified in aphid genomes also originated from HGT events, we conducted a phylogenetic analysis to examine the evolutionary relationships with sequences from 288 insects (Supplementary table S2), 36 plants, 32 bacteria, and 31 fungi (Supplementary table S3).

The protein sequences of upstream genes (*GPS*, *PS*, and *CD*) in insects were clustered closely with fungal sequences but remained distinct from plant and bacterial sequences (Fig. 2A). In contrast, the protein sequences of *CCO* in insects formed a distinct cluster (Fig. 2A), separated from the sequences of fungi, plants, and bacteria, suggesting *CCO* are likely native insect genes instead of horizontally transferred genes.

**Fig. 2.**
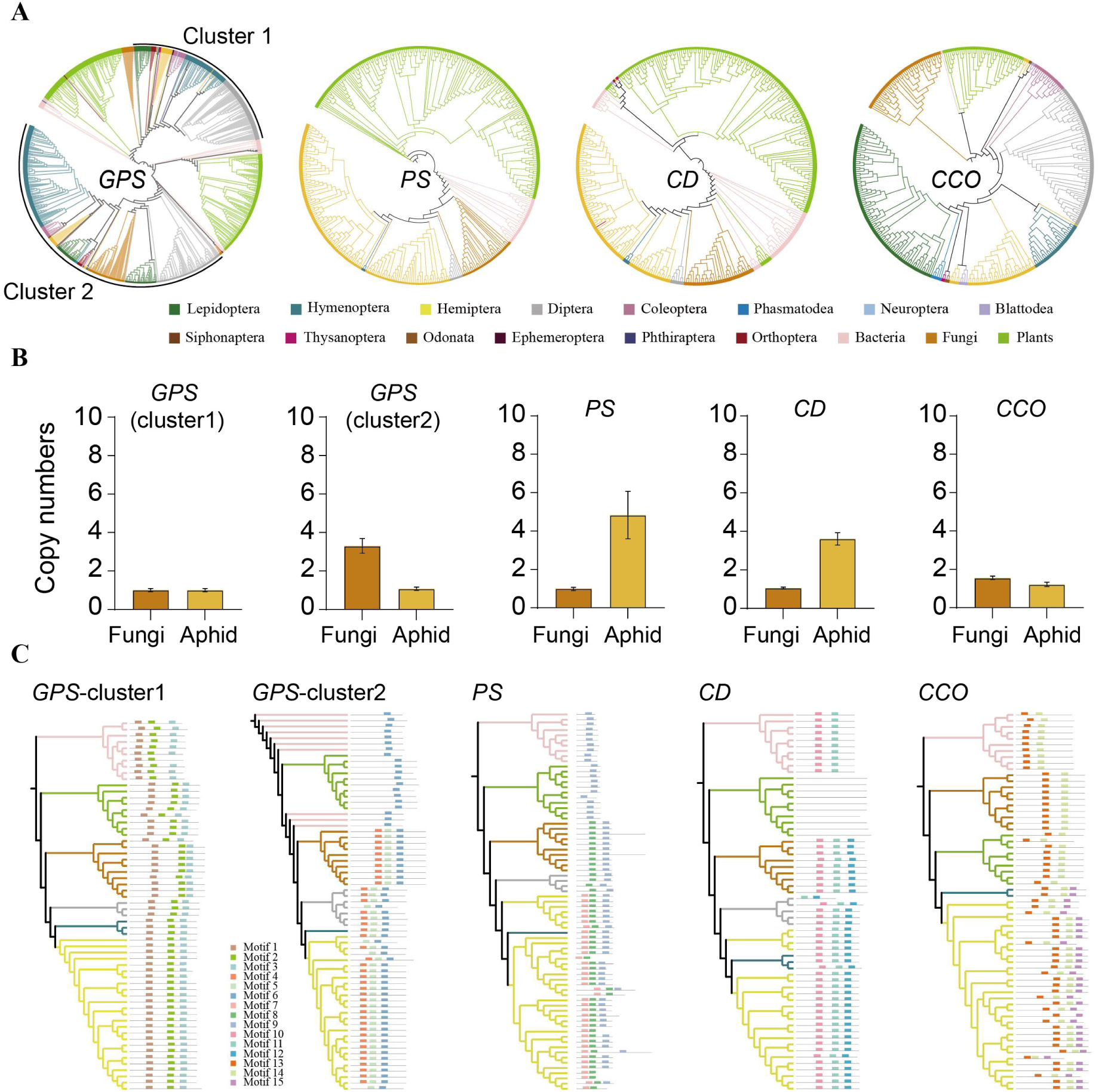
*GPS*, *PS*, and *CD* genes are horizontally transferred from fungi, while *CCO* is of insect origin. (A) ML trees illustrating the evolutionary relationships between insects and species from three biological kingdoms. Trees are rooted at the midpoint. Colored branches classify species into distinct kingdoms and insect orders. The color-based groups are shown below the trees. The *GPS* tree contains two clusters: the upper insect cluster is designated as Cluster 1, while the lower cluster is Cluster 2. (B) Average gene copy numbers in fungi and aphids. *PS* and *CD* genes exhibit higher copy numbers in aphids than in fungi. The y-axis represents the average copy number per species (total copy numbers/total species). The total gene copy number is calculated based on the total protein sequences in each species. Fungal and aphid species included correspond to those analyzed in (A). (C) Motif analysis reveals identical motif profiles between insects and fungi. *GPS* cluster1 is conserved across different species, while *CCO* is specific to insects. Only the top 10 protein sequences from fungi, bacteria, and plants were included in these trees. Insect protein sequences come from those species containing *GPS*, *PS*, *CD*, and *CCO* genes simultaneously. Branch colors are consistent with (A). Colored rectangles indicate distinct motif features, with lines below motifs representing relative protein sequence lengths.

Sequences of aphid *tor* genes in the tree of *CD* were clustered with fungal sequences, verifying the results of the blast approach (Fig. 2A). *GPS* had two distinct clusters, and both were grouped with fungi (Fig. 2A). In *GPS* cluster 1, fungal sequences formed the outer group, while insect sequences formed the inner group, a pattern also observed for *PS* and *CD* (Fig. 2A). It indicated that the HGT of those genes likely occurred in ancestor insects. Interestingly, in cluster 2, *GPS* sequences from fungi appeared embedded with sequences from Lepidopteran species, suggesting this copy of *GPS* may have originated from a separate HGT event.

Except for the aphid species in Hemiptera, many insect species in other orders did not have *PS* and *CD* genes, suggesting the acquired genes may be lost during speciation (Supplementary fig. S2C and D). The number of homologs in aphid species was higher than in other insect species and fungi (Fig. 2B, Supplementary fig. S2 C and D). These genes were single-copy in most fungal species, while they were multiple copies in aphid species (Fig. 2B), indicating these genes have been duplicated in aphid species after the original copy was acquired from fungi by the ancestor species.

Motif analysis revealed that insect sequences had similar motif profiles as fungal sequences and differed from those of plants and bacteria (Fig. 2C), implying that those genes have been horizontally transferred from fungi. Additionally, the distinct motif patterns between *GPS* cluster 1 and cluster 2 suggested that these two clusters may have originated from independent HGT events (Fig. 2C). Hemipteran *PS* sequences contain an additional motif absent in dipteran insects and fungi sequences (Fig. 2C), suggesting potential functional diversification after gene acquisition.

### Carotenogenic genes are clustered in the chromosome and expressed in different aphids with distinct colors

We selected four aphid species with distinct color patterns to further investigate carotenogenic genes. *Brevicoryne brassicae* is greyish, *Rhopalosiphum padi* is black, while *Acyrthosiphon pisum* and *M. persicae* display a remarkable range of color polymorphisms, varying from green to reddish depending on the clones (Fig. 3A).

**Fig. 3.**
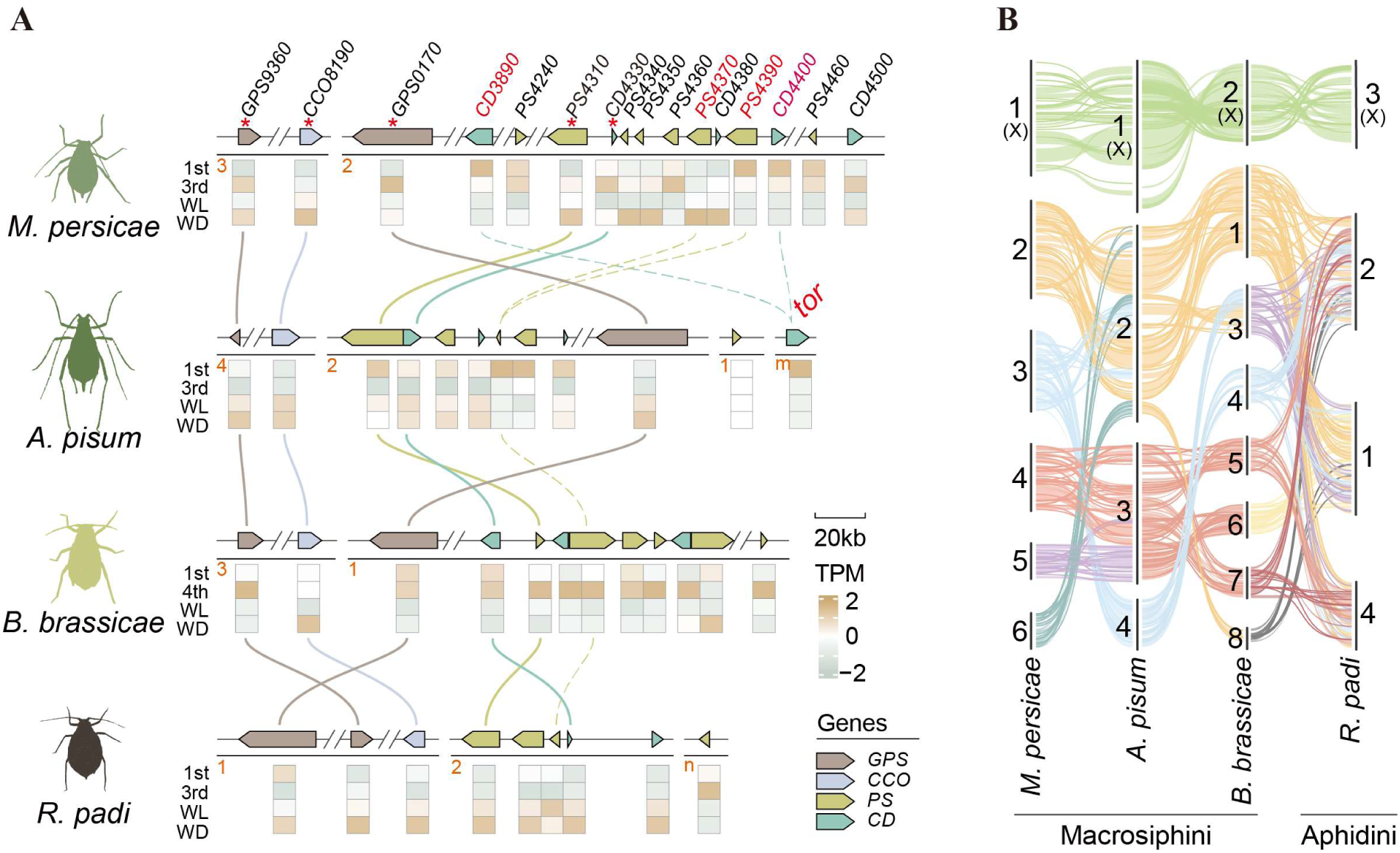
Tandem arrangement and expression dynamics of carotenogenic genes in aphid species. (A) A schematic diagram illustrates the morphology and body coloration of four aphid species. In the gene structure diagram, black lines under the arrows represent chromosome length. The red numerical values correspond to scaffolds. m: NW_021771109.1; n: scaffold_116. The "//" symbol indicates a folded distance between genes. Gaps separate different chromosomes. Arrows denote gene orientation, with arrow colors corresponding to different genes. Gene IDs marked in red indicate duplicate genes in *M. persicae*. Dashed lines connect the genes clustered with these duplicate genes. Red stars highlight collinear genes among the four aphid species. Collinear genes are linked by solid curves. The heatmap displays transcriptomic TPM values across development stages (the first instar and third instar) and morphs (wingless (WL) and winged (WD) forms). *B. brassicae* is shown at the fourth instar instead of the third. (B) Syntenic analysis among the four aphid genomes reveals differences between the Macrosiphini and Aphidini lineages. Colored chains represent syntenic blocks. Arabic numerals correspond to scaffolds in *M. persicae*, *B. brassicae*, and *R. padi*. The X chromosome for each species is located on the topset. In *A. pisum*, numerals 1 to 4 correspond to NC_042493.1, NC_042494.1, NC_042495.1, and NC_042496.1, respectively.

Comparative genomic analysis revealed the clustering of carotenogenic genes in aphid genomes. All four species harbored two copies of *GPS*, with one copy syntenic to *CCO* gene (Fig. 3A). In *R. padi* (Aphidini lineage), the second *GPS* copy was colocalized on the same chromosome as the first. In the Macrosiphini species (*B. brassicae*, *A. pisum*, and *M. persicae*), this copy was located on a separate chromosome and clustered with *PS* and *CD* genes (Fig. 3A). This pattern may have arisen from extensive autosomal rearrangements during aphid speciation (Fig. 3B). *M. persicae* had the highest number of *PS* (8 copies) and *CD* (5 copies) genes, whereas *B. brassicae* (6 *PS*, 3 *CD*), *A. pisum* (5 *PS*, 4 *CD*), and *R. padi* (4 *PS*, 2 *CD*) harbored fewer copies. In *M. persicae*, *PS* and *CD* genes were tandemly duplicated within the same chromosome (Fig. 3A, Supplementary fig. S3).

Notably, a homolog of *tor* (*CD*) in *A. pisum* was present as two copies in *M. persicae* but was absent in the other two species (Fig. 3A, Supplementary fig. S4A). These two copies were part of a duplicated gene cluster unique to *M. persicae* (Supplementary fig. S4A). In *M. persicae*, *PS* genes underwent multiple duplicate events. For example, *M. persicae* contained two copies of *PS* genes (*PS4390* and *PS4370*), while only one copy was found in the other aphid genomes (Supplementary fig. S4B). A similar duplication pattern was observed for *PS4340* and *PS4350* (Supplementary fig. S4B). These findings suggest that carotenogenic genes have undergone multiple gene gain-loss events during aphid diversification.

Carotenoids are key pigments contributing to insect body coloration, with aphids displaying extensive variation, likely reflecting species-specific differences in carotenoid biosynthesis. To explore these differences, we analyzed the expression of identified carotenogenic genes across the four species using publicly available RNA-seq data (Fig. 3A). Despite differences in gene copy numbers, most carotenogenic genes were transcriptionally active, except *PS-1759* in *A. pisum* (Fig. 3A). Gene expression levels varied across developmental stages, indicating dynamic regulation of the carotenoid biosynthetic pathway in aphids. Notably, several duplicated gene copies exhibited morph-specific expression patterns. In *M. persicae*, multiple duplicated copies of *PS* (*PS4310*, *PS4340*, *PS4350*, *PS4370*) and *CD* (*CD4380*, *CD4500*) were more highly expressed in winged morphs compared to wingless ones. Similar expression patterns were observed in the other three species, suggesting that duplicated copies may have undergone functional diversification, potentially contributing to morph- or stage-specific carotenoid production.

### A *PS* gene is a novel component that likely contributed to the red color

To assess the contribution of identified carotenogenic genes in the aphid coloration, we compared the expression levels of these genes in *M. persicae* clones with distinct body colors, the greenish clone and the reddish clone (Fig. 4A). *GPS* genes, and most *PS,* and *CD* genes showed no significant difference between the two clones, indicating they were unlikely to play critical roles in the aphid coloration (Fig. 4B-G). *CD-4400* and *CD-3890*, two homologs of the pea aphid *tor* gene, exhibited opposite expression patterns between greenish and reddish clones. *CD-4400* was expressed 10.84 times higher in the reddish clone than in the greenish clone, while *CD-3890* was 2.71 times higher in the greenish clone (Fig. 4F and G). *CD-4400* has been reported that its expression correlate with the red color in *M. persicae*, but *CD-3890* is not (Ge et al. 2024). Similar to *CD-3890*, genes such as *PS-4350*, *CD-4380*, and *CCO* were highly expressed in the greenish clone compared to those in the reddish clone (Fig. 4D, G, and H), suggesting they were unlikely to contribute to the red color. Interestingly, similar to *CD-4400*, the expression of *PS-4390* was 15.31 times higher in the reddish clone compared to that in the greenish clone (Fig. 4E). This indicated that *PS-4390* may also be a key gene involved in aphid red color formation.

**Fig. 4.**
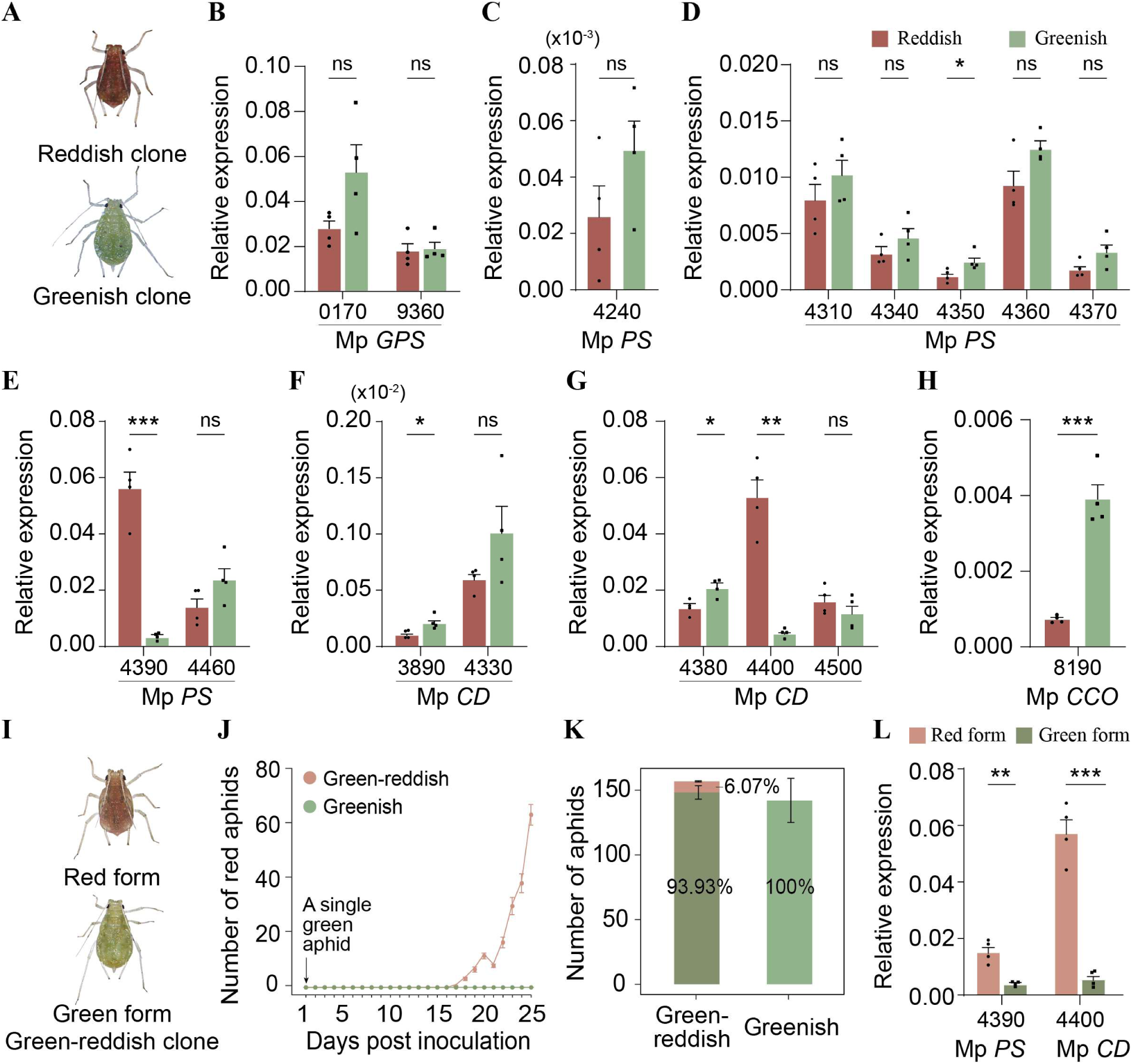
Carotenogenic genes exhibit diverse expression patterns in red and green color polymorphisms, with *PS-4390* and *CD-4400* significantly upregulated in red-colored aphids. (A) Images of reddish and greenish clones of *M. persicae*. (B-H) Gene expression comparisons between reddish and greenish clones. Relative expression levels were calculated as 2^-ΔCt relative to *RPL7*. The reddish clone is represented using red-brown, while the greenish clone is shown in grayish green. Solid squares indicate replicates. Y-axis values are scaled using (×10⁻²) or (×10⁻³) for better visualization of small values. (I) Images of red and green aphids from the green-reddish clone of *M. persicae*. (J) The number of red aphids increased over time in the green-reddish clone, while the greenish clone served as a control. A single green aphid was inoculated onto the host plant, and red offspring were counted over time. (K) Total aphid counts in the population. Percentages indicate the proportion of each color within the total population. (L) Relative expression levels of *PS-4390* and *CD-4400* in the green-reddish clone. The red form of the green-reddish clone is shown in light red-orange, and the green form is displayed in dark grayish green. ns: not significant; *: *p*<0.05; **: *p*<0.01; ***: *p*<0.001. All pairwise comparisons were performed using an unpaired two-tailed *t*-test.

The greenish and reddish clones are different aphid lineages with genetic variations. The differences in the expression of *PS-4390* and *CD-4400* may not be related to color but the results from genetic variations in regulatory elements. Another *M. persicae* clone, which was named a green-reddish clone here, exhibits green and red color polymorphism (Fig. 4I). A single parthenogenetic green nymph was placed on the host plant, and the red aphids appeared about 16 days later (Fig. 4J). The red aphids were counted at approximately 6.07% of the total population (Fig. 4K). To assess the impact of *PS-4390* and *CD-4400* on red color, we compared the expressions of these two genes in green and red form. Expressions of *PS-4390* and *CD-4400* were 3.84 times and 9.87 times higher in red forms than in green forms, respectively (Fig. 4L). The green and red forms were genetically identical, the differences in expressions of *PS-4390* and *CD-4400* were highly related to red color formation. *PS-4390* was likely a novel component that contributed to red formation in aphids.

### Knocking down the expression of *PS-4390* causes the fading of red coloration in red aphids

To investigate the involvement of *PS-4390* in red coloration, we knocked down the expressions of *PS-4390* in both the reddish clone and the red aphids of the green-reddish clone (Fig. 5A). In addition, we also knocked down the expression of *CD-4400* in these two kinds of red aphids (Fig. 5A). As expected, knocking down the expression of *CD-4400* resulted in a loss of redness in both reddish clones and red aphids of the green-reddish clone (Fig. 5A). Similarly, silencing *PS-4390* led to a comparable loss of red coloration (Fig. 5A), increasing green forms in the population by approximately 11.25% and 7.5% of RNAi-treated aphids in the two clones, respectively (Fig. 5B). In both RNAi experiments, aphids that turned green exhibited significantly higher RNAi efficiency compared to controls (Fig. 5C, D). The comparable proportions of red aphids losing pigmentation upon *PS-4390* and *CD-4400* knockdown support the involvement of both genes in the same biosynthetic pathway.

**Fig. 5.**
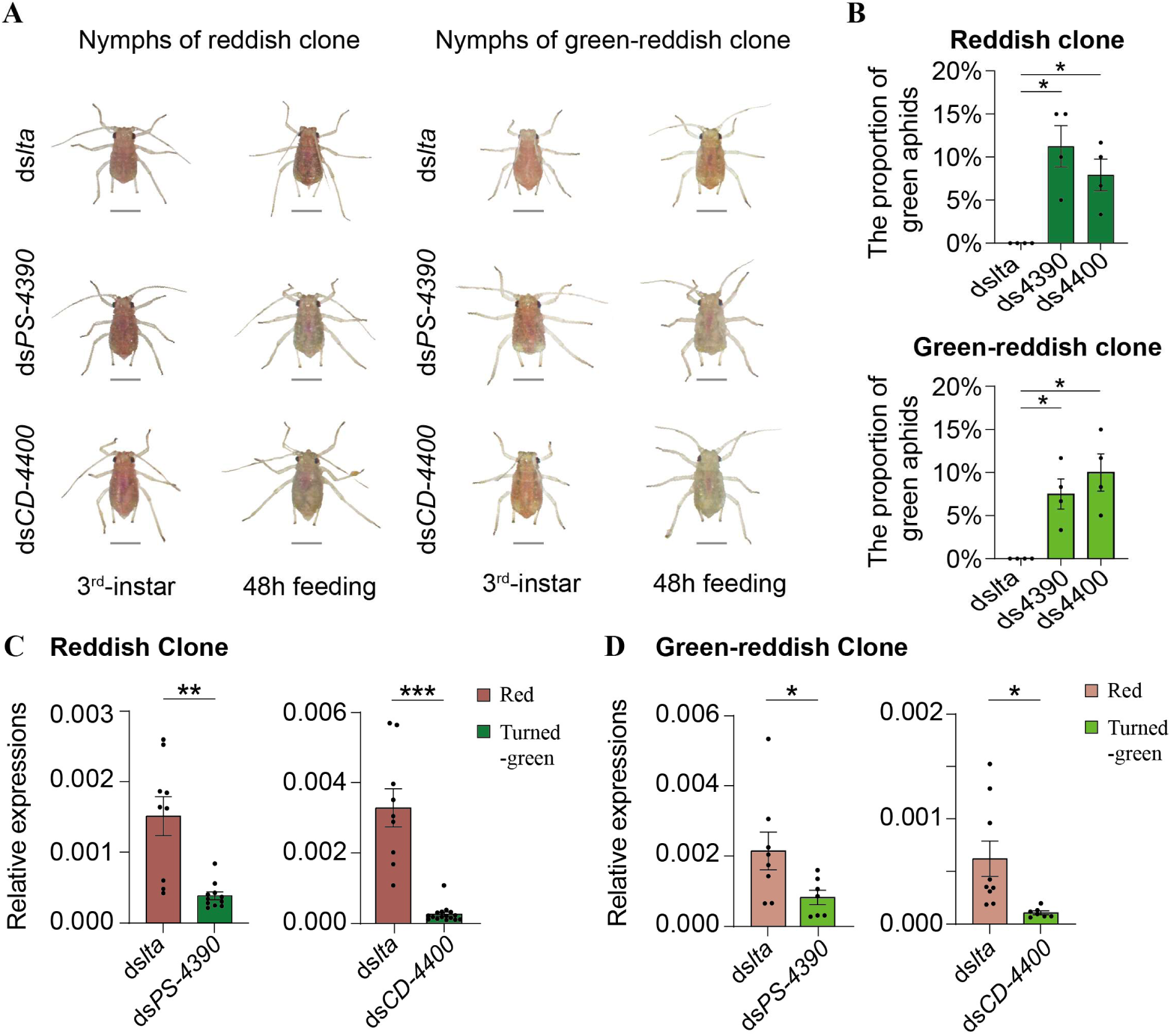
Silencing *PS-4390* and *CD-4400* caused red aphids to fade to green. (A) Comparison of aphid body color before and after 48 hours of dsRNA feeding in red aphids. Aphids were fed with dsRNA at the third instar stage. All scale bars: 500 µm. (B) The proportion of green aphid following dsRNA feeding. Green aphids represent those that faded from red. The number of green aphids was compared to the control (ds*lta*) using *t*-test, with *p* < 0.05 indicating a significant effect of dsRNA feeding on the loss of red coloration. *: *p*<0.05; **: *p*<0.01; ***: *p*<0.001. c-d) Gene expression levels after RNAi in the reddish clone (C) and green-reddish clone (D). Relative expression levels were calculated as 2^-ΔCt relative to *EF-1α*. Solid circles indicate replicates. ns: not significant. All pairwise comparisons were performed using an unpaired two-tailed *t*-test.

## Discussion

Horizontal gene transfer (HGT) is a widely observed phenomenon and is considered a driving force for adaptation and evolution (Schönknecht et al. 2014). In insects, HGT is more common than previously believed (Li et al. 2022). So far, studies of HGTs in insects are focused on the single events that are incorporated into existing pathways (Lapadula et al. 2020; Xia et al. 2021). HGTs for genes involved in a complex multi-step pathway is rarely reported. To address this question, we examined the carotenoid biosynthesis pathway as a model for understanding the HGT of complex pathways in insects. We analyzed 26 genes previously reported to participate in carotenoid biosynthesis in plants, fungi, and bacteria and searched for homologous sequences in 23 aphid genomes. Our analysis revealed that *GPS*, *PS*, *CD*, and *CCO* are widely present in aphid genomes as well as in the genomes of other insects. Phylogenetic analysis showed that *GPS*, *PS*, and *CD* were acquired by insects via HGT from fungi, while *CCO* was identified as a native insect gene. Notably, most insect genomes contain two copies of *GPS,* and these two copies have different motif profiles likely resulting from independent HGT events. In aphid genomes, *PS* and *CD* have undergone extensive gene duplication, a pattern not observed in many other insects. We further assessed the expression of genes in the carotenoid biosynthesis pathway across aphid species with distinct body colors by analyzing the expression patterns of the homologous genes. Despite variations in body color, these genes were expressed across different aphid species, showing substantial variability in expression levels. Particularly, we compared gene expression between clones of *M. persicae* with different colors, including reddish and greenish clones, as well as different color forms within a single clone (green-reddish clone). Our results revealed that *CD-4400*, a homolog of the pea aphid *tor* gene, is associated with redness. More importantly, we identified *PS-4390* as a novel component contributing to redness in *M. persicae*. These findings highlight the complexity of HGT events that can incorporate multiple genes of the same pathway into the genomes of recipient species.

In *A. pisum*, three carotenoid synthase/cyclase genes and four carotenoid desaturase genes were first reported by Moran and Jarvik (2010). Subsequent work expanded the synthase/cyclase gene count to four (Takemura et al. 2021; Ding et al. 2022), although the specific intermediates and functional roles of individual gene copies remain unresolved. A recent study reported 11 such genes in *M. persicae* (Trissi et al. 2023), while our analysis detected eight, potentially reflecting differences in genome assembly quality. Notably, expression patterns of *CD-4400* and *PS-4390* (also reported as *CDE-2* and *CCS-9* in Trissi et al. 2023, respectively) were elevated in red clones, corroborating our findings. *CarD763*, recently implicated in regulating red pigmentation (Ge et al. 2024), also corresponds to *CD-4400*, further supporting the robustness of our gene annotation.

Genes at downstream of the carotenoid biosynthesis pathway, such as *CCO* gene, are already present in insect genomes. However, several upstream genes in this pathway were acquired by ancestral species through HGT from fungi. In fungi, *PS* and *CD* genes are often organized in clusters on the same chromosome (Supplementary fig. S3, Avalos et al. 2017). Similarly, we observed the clustering of *PS* and *CD* genes in aphid chromosomes (Supplementary fig. S3). Since the original fungal gene copies are likely to have been lost during speciation events, it remains uncertain whether the genes required for this pathway were transferred to insects simultaneously or sequentially. Genes such as *PS*, and *CD* have also been identified in certain *Tetranychus* species (Grbić et al. 2011, Bryon et al. 2017), suggesting that the HGT of these genes may have occurred in ancestral arthropods. It is unclear whether the copies found in aphid genomes resulted from independent HGT events or are descendants of genes acquired by ancestral arthropods. *PS* and *CD* genes were also found in the genomes of *Adelges cooleyi* and *Daktulosphaira vitifoliae* but were lacking in most other Hemipteran species, such as *B. tabaci*, *Nilaparvata lugens* (Supplementary table S2). This implies that these genes are more likely derived from the common ancestor of Adelgidae and Phylloxeridae. *PS* and *CD* are typically single-copy genes in most fungal species (Supplementary fig. S2), whereas they exist as multiple copies in the genomes of Adelgidae, Phylloxeridae, and Aphididae (Supplementary fig. S2). The pattern suggests that these genes have undergone extensive duplication in these species following the loss of the original HGT-derived copies.

Many insect species are believed to lack the ability to synthesize carotenoid compounds and instead rely on acquiring them from their diet (Clark and Lampert 2018). Here, we discovered that many aphid genomes contain a complete carotenoid synthesis pathway. This indicates that aphids are capable of *de novo* synthesizing carotenoids. In addition to the widely conserved horizontally transferred gene (*GPS*) and the native gene *CCO*, two key genes *PS* and *CD* were found in aphid genomes which are absent in the vast majority of insect genomes (Fig. 1A, Supplementary table S2). Interestingly, some insect species, such as the dipteran species *Bradysia coprophila* and *Sitodiplosis mosellana* and the hymenopteran species *Aphidius gifuensis*, also possess at least one copy of both *PS* and *CD* genes (Supplementary table S2). In *S. mosellana*, carotenogenic genes have been previously identified and are hypothesized to contribute to ecological diversification across host lineages (Cobbs et al., 2013). This suggests that the entire carotenoid synthesis pathway has been evolutionarily selected to be maintained in some insect species, with a notably higher prevalence in aphids.

We observed that certain differentially expressed duplicated copies of *PS* and *CD* genes were not associated with coloration in *M. persicae*. For example, *CD-4380* and *CCO-8190* were highly expressed in the greenish clone compared with the reddish clone (Fig. 4G and H), while *PS-4360* and *PS-4460* were highly expressed in the red forms of the green-reddish clone compared with the green forms (Supplementary fig. S5C). Additionally, several duplicated genes showed morph- or stage-specific expression patterns (Fig. 3A), indicating potential functional divergence. This pattern of gene duplication and functional diversification is consistent with the concept of neofunctionalization, where duplicated copies of HGTs may acquire novel functions while one copy retains its original role. For instance, in some lepidopteran pests (*Plutella xylostella* and *Helicoverpa armigera*), horizontally acquired bacterial genes confer resistance to *Bacillus thuringiensis* (Bt) toxins (Jurat-Fuentes et al. 2021). Gene duplication allowed one copy to maintain its ancestral function while the other evolved increased specificity or broader resistance mechanisms. Similarly, in beetles like *Tribolium castaneum*, horizontally transferred bacterial chitinase genes, which break down chitin, underwent duplication. The duplicated copies then diversified into gut-specific and systemic enzymes, enhancing digestion and possibly contributing to antifungal defense (Wang et al. 2023).

In summary, HGTs have also occurred for genes involved in complex multi-step pathways, although the precise trajectory of horizontal transfer for each component of these pathways remains unclear. After their initial acquisition, HGTs undergo extensive duplication in some insect species, and some duplicated copies may acquire novel functions.

## Materials and Methods

### Aphid clones and rearing

Three *M. persicae* clones are involved in this study. According to their diverse phenotypes in body color, we distinguished the three clones. The reddish clone is a red population, the greenish clone is green, and the green-reddish clone exhibits green and red color polymorphism. All clones maintain their phenotypes under laboratory rearing conditions. The reddish clone was originally collected from a tobacco plant and was obtained from the laboratory of Prof. Feng Ge (Institute of Zoology, Chinese Academy of Sciences, Beijing, China). The greenish clone was sourced from the laboratory of Prof. Tongxian Liu (Wu et al. 2024), while the green-reddish clone was provided by Prof. Jianqiang Wu (Kunming Institute of Botany, Chinese Academy of Sciences, Kunming, Yunnan, China).

All aphid stocks were maintained on individual seedlings enclosed in plastic cages at 24 ± 1 °C, under a 16/8 h light/dark photoperiod, with 60 ± 10% relative humidity. The reddish clone was reared on *Nicotiana benthamiana*, while the other two clones were reared on *Arabidopsis thaliana* (Col-0). All aphid species have been maintained in our laboratory for more than three years.

### Carotenogenic genes in bacteria, fungi, and plants

Eleven carotenogenic genes from plants (Badejo 2018; Stanley and Yuan 2019; Stra et al 2023), seven from bacteria (Giraud et al. 2004; Klassen 2009), and eight from fungi (Avalos et al. 2017) were compiled from published literature. The protein sequences corresponding to these genes were retrieved from the UniProt database using gene names and source species as keywords. *A. thaliana* was selected as the source plant species. *Fusarium fujikuroi* was chosen as the fungal representative based on the literature (Avalos et al. 2017). Bacterial source species were selected from a diverse set of photosynthetic bacteria, following previous studies (Giraud et al 2004; Klassen 2009). The collected protein sequences were assembled into a custom query set.

Organisms from different phyla across the plant, bacteria and fungi kingdoms were included, to ensure comprehensive taxonomic representation. These organisms were required to possess complete annotated genomes available in the NCBI genome database (accessed on March 8, 2024). Additionally, the top 20 organisms for each gene, identified through online NCBI BLASTp, were included. Protein FASTA files were directly downloaded from NCBI RefSeq and used to construct local BLAST databases via makeblastdb command in BLAST 2.5.0+ (Camacho et al. 2009). Local BLASTp searches were then conducted to identify homologs of the 26 carotenogenic genes across plant, bacterial, and fungal species, using an e-value threshold of <1e-15 (Xia et al. 2021) and a minimum sequence length of 100 amino acids. (Sakurai et al. 2007).

### Insect genomes

A total of 288 insect species from 14 orders were selected for this study (Supplementary table S2). Of the 288 insect species, 23 are aphids, whose genomes were obtained from public databases (NCBI and BIPAA) and published data (Mathers et al 2022; Liu et al. 2024a). Protein sequences were extracted from the genomes using GffRead v0.12.7 (Pertea and Pertea 2020), and a custom aphid database was constructed using the makeblastdb command in BLAST 2.5.0+.

The remaining 265 insect species were selected from the NCBI genome database using the keyword “Insecta” and filtered based on RefSeq annotation. Protein FASTA files for these species were retrieved directly from the NCBI genome, and used to construct a custom insect database in BLAST 2.5.0+. Local BLASTp searches were performed against both the aphid and insect databases using the custom query set, with filtering criteria set at an e-value < 1e-15 and sequence length > 100 amino acids.

### Pfam analysis

Homologous sequences obtained from BLASTp were subjected to Pfam domain annotation using HMMER v3.4 with the pfam_scan v1.6 tool (Eddy 2011), setting an e-value threshold of 1e-5. Based on similarity in gene families and identical Pfam domains, the 26 carotenogenic genes were ultimately consolidated into 15 genes. These genes include: *GPS* genes (polyprenyl_synt domain, PF00348.21), *PS* genes (SQS_PSY domain, PF00494.23), *CD* genes (Amino_oxidase domain, PF01593.28), *LC* genes (Lycopene_cycl domain, PF05834.16), *CarT* genes (RPE65 domain, PF03055.19), *CarS* genes (LON_substr_bdg domain, PF02190.20), *CarD* genes (Aldedh domain, PF00171.26), *ZCIS* genes (NnrU domain, PF07298.15), *β-OHase* genes (FA_hydroxylase domain, PF04116.17), *ZEP* genes (FHA domain, PF00498.30), *crtA* genes (no known Pfam domain), *crtF* genes (Methyltransf_2 domain, PF00891.22), *crtC* genes (no known Pfam domain), *CarO* genes (Bac_rhodopsin domain, PF01036.22), *CarX* genes (RPE65 domain, PF03055.19).

### Phylogenetic analysis

Protein sequence alignments were conducted using ClustalW v2.1 (Thompson et al. 2002), and ambiguously aligned regions were trimmed with trimAl v1.4 (Capella-Gutiérrez et al. 2009) using the “-automated1” parameter. Maximum likelihood (ML) trees were constructed with IQ-TREE v2.2.2.7 (Nguyen et al. 2015), applying the “-m TEST” option to automatically select the best-fit model of amino acid substitution and “-B 1000” to conduct ultrafast bootstrap analysis (Minh et al. 2013; Li et al. 2022). The best-fit models for each gene tree were selected as follows: LG+G4 for *GPS*, JTT+G4 for *PS*, Q. pfam+G4 for *CD* and Q. insect+I+G4 for *CCO*. Newick tree files were rooted at the midpoint using the ape R package (Paradis and Schliep 2019).

Phylogenetic trees for motif visualization were constructed using RaxML v8.2.13 (Stamatakis 2014) with the “-f a -# 1000” option to rapidly compute 1000 Bootstrap iterations, “-m PROTGAMMALGX” to specify the amino acid substitution model, and “-o” to designate bacteria as the outgroup. For these trees, only the top 10 protein sequences from plants, bacteria, and fungi were selected, along with insect species from three orders: Hemiptera, Diptera, and Hymenoptera. Finally, tree files were uploaded to the iTOL v5 (Letunic and Bork 2021) for visualization.

### Motif analysis

Protein FASTA files used in RaxML were uploaded to the MEME Suite 5.5.7 website (Bailey et al. 2015) for motif identification, and three motifs were automatically generated. The MEME XML outputs were downloaded and imported into the iTOL website to create MEME datasets, which was subsequently used to add tree colors and annotations in Adobe Illustrator 2022.

### Genome syntenic analysis

Genome information used in syntenic analysis is listed in Supplemental table S3. The genome of *A. pisum* was used the latest version downloaded from AphidBase (International Aphid Genomics Consortium, 2018). Gene structure information, including start and end positions, orientations, and chromosomal locations of carotenogenic genes, was extracted from the aphid GFF annotation files.

The fungal genomes mentioned in Supplementary fig. S3 were from two species, *F. fujikuroi* (designated as the source species in the previous analysis) and *Podospora anserina* (selected due to its high diversity of carotenogenic genes). Gene structure diagrams were drawn using the ChiPlot website (https://www.chiplot.online), where tables containing gene structure information and gene name annotations were uploaded for automated visualization. Additionally, breaks were set to fold long genomic distances between genes.

Genome syntenic analysis was performed using TBtools-II (Chen et al. 2023). The Genome Length Filter tool of TBtools-II was employed to remove short contigs, generating updated genome FASTA and annotation files. The One Step MCScanX tool was used to detect gene synteny and collinearity. Finally, the Dual Synteny Plot tool was applied to extract collinear blocks of carotenogenic genes between two species.

### Transcriptomic analysis

All sequencing reads involved in the transcriptomic analysis were downloaded from a public database (NCBI SRA), with detailed information provided in Supplementary Table S5. The reference genomes of the four aphid species were consistent with those used in the syntenic analysis.

Sequencing reads were aligned to the reference genomes using RMTA v2.6.4 (Peri et al. 2020), with default parameters set as ‘-m 20 -f 1 -k 1”, filtering intron length (-m), base coverage (-f) and FPKM threshold (-k). Additionally, Hisat2 mapping, FastQC quality control, and duplicates removal were performed using the “-t -e -d” options integrated in RMTA (Andrews 2010; Langmead and Salzberg 2012; Kim et al. 2019).

Feature counting was conducted using HTseq v2.0.8 with the “-t gene” option to specify gene-level features (Putri et al. 2022). TPM values were calculated using TPMCalculator v0.0.5 with the “-e” option to obtain transcript-level expression values (Vera Alvarez et al. 2019).

### RNA extraction

Aphids of *M. persicae* were collected by flash freezing in liquid nitrogen, with 10 apterous fifth-instar nymphs per tube. For RNAi efficiency detection in *M. persicae*, a single aphid was collected per tube. Each sample was ground to a fine powder using the High-Speed Tissue Grinder JXFSTPRP-64L (Jingxin, Shanghai, China).

Samples for RNA extraction were thoroughly dissolved in TRIzol® Reagent (Invitrogen) following the manufacturer’s instructions. RNA concentration and purity were measured using a Microspectrophotometer (K5600C, KAIAO, Beijing, China). High-quality RNA was transcribed into complementary DNA (cDNA) using 1μg of total RNA with a HiScript II 1st Strand cDNA Synthesis Kit (Vazyme, Nanjing, China) according to the manufacturer’s instructions.

### qRT-PCR

Quantitative detection was performed using the CFX Connect™ Optics Module Real-Time System (Bio-Rad) with the Hieff® qPCR SYBR Green Master Mix (No Rox) (Yeason, China). Each reaction was carried out in a total volume of 20 µL, consisting of 10 µL SYBR Green Mix, 0.4 µL of each primer, 2 µL cDNA, and 7.2 µL nuclease-free water (Invitrogen). Amplification conditions were as follows: 95°C for 5 minutes, followed by 40 cycles of 95°C for 10 seconds and 60°C for 30 seconds. The specificity of qRT-PCR primers was further confirmed by a melting curve analysis.

Gene expression levels were calculated using the 2^-ΔCt^ method (Yuan et al. 2006), with *ribosomal protein L7* (*RPL7*) and the *nuclear elongation factor 1-alpha (EF1α)* used as reference genes. The *RPL7* gene was used in the experiments to detecting gene expression levels, and the *EF1α* gene was used in the detection of RNAi efficiency.

Primers were designed using Beacon Designer 7 and NCBI Primer-BLAST (Supplementary table S4, Ye et al. 2012), followed by fine-tuning in SnapGene® 5.2 to ensure complete gene specificity. Three pairs of primers were initially tested, and their specificity was assessed based on melting-curve analysis, with only one melting-temperature (Tm) peak selected. Additionally, electrophoresis was performed to verify the presence of a single band. The most specific primer pair was selected for further study.

### RNA interference

Double-stranded RNA (dsRNA) fragments (300-500 bp) were designed for target gene silencing. These fragments were synthesized using the T7 High Yield RNA Transcription kit (Vazyme, Nanjing, China) and purified with a phenol: chloroform: isoamyl: alcohol (25:24:1) solution, following the manufacturer’s protocol. The *Lymphotoxin A* (*LTA*) gene from a mouse (Bai et al. 2022) was synthesized as a control dsRNA.

Purified dsRNA was dissolved in nuclease-free water, and its concentration was measured using a KAIAO Microspectrophotometer. dsRNA at a final concentration of 1.2 μg/μl was incorporated into an artificial diet, containing 30% sucrose and 0.02% methyl red, with a total volume of 50 microliters (Bilgi et al. 2017). Fifteen third-instar aphids were transferred into cylindrical containers (25 mm in diameter × 15 mm in height) for dsRNA feeding.

After 48 hours of continuous feeding, a single aphid was collected per tube for RNA extraction to assess RNAi efficiency, with more than nine replicates per experiment. Phenotypic changes were recorded by photographing aphids that turned green, while those remaining red were individually collected and analyzed for RNAi efficiency. Each experiment included at least seven replicates.

### Statistical analysis

Bioinformatics data were generated using Linux-based pipelines, and data processing was performed using Python scripts. All graphical visualizations were produced using R packages. Pairwise comparisons were conducted using two-tailed *t*-tests.

## Supplementary Material

Supplementary material is available online.

## Acknowledgments

The authors thank Prof. Tongxian Liu (Guizhou University, China), Prof. Jianqiang Wu (Kunming Institute of Botany, CAS, China), Prof. Feng Ge (Institute of Zoology, CAS, Beijing, China) for kindly providing aphid colonies. We also appreciate Yongjian Liu and Gangqi Fang for their guidance in bioinformatic analysis.

## Data Availability

All study data are included in the article and supplementary material.

## Author contribution

Y.C. conceived and designed the study. R.H. conducted the majority of bioinformatic analyses. R.H., J.W., S.L. and P.Y. performed the experiments. Y.C. and R.H. drafted the manuscript. S.Z., G.W., and C.N. provided academic consultation and contributed to manuscript revision.

---

**Supplementary fig. S1.**
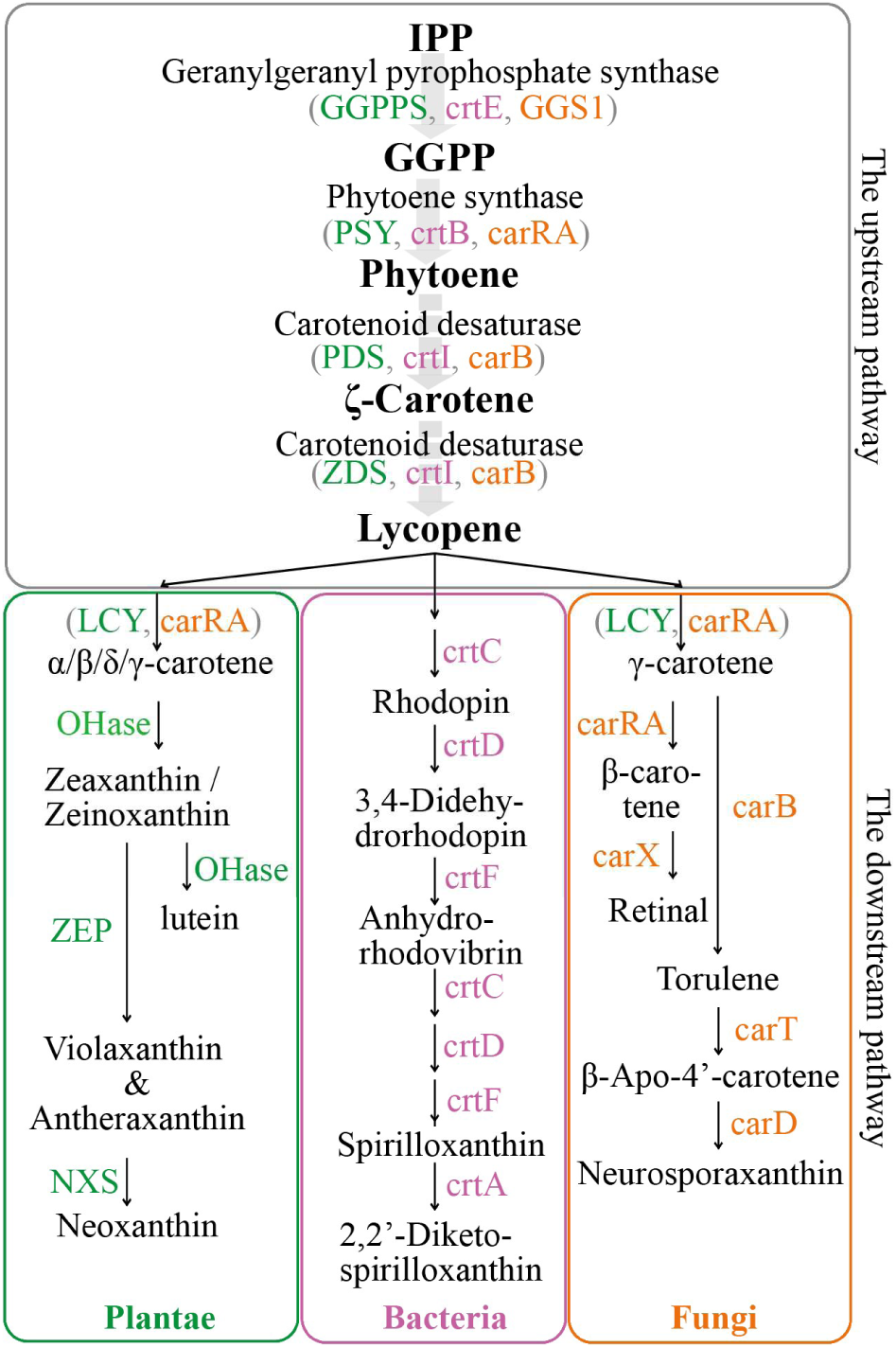
Schematic representation of carotenoid biosynthesis pathway across plant, bacteria, and fungi kingdoms. Enzyme abbreviations are color-coded: green for plantae, pink for bacteria, and orange for fungi. Conserved pathways across the three kingdoms (the upstream pathway) are depicted with gray wide arrows, while dashed wide arrows indicate folding steps. Thin solid arrows represent kingdom-specific pathways (the downstream pathway). The upstream pathway products are highlighted in bold font, while the full names of enzymes are shown in regular font. The products of kingdom-specific genes in the downstream pathway are also in regular font.

**Supplementary fig. S2.**
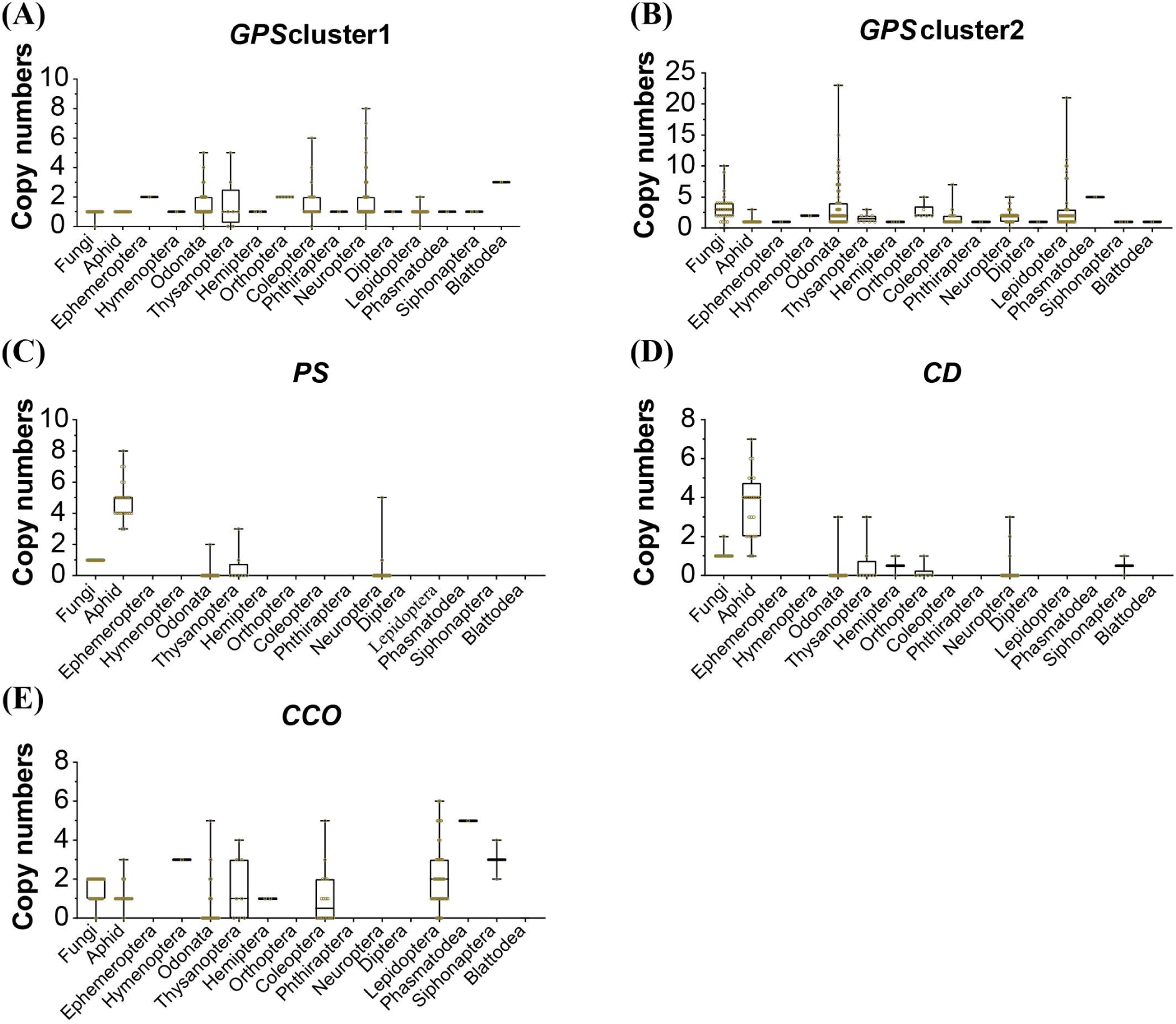
Average copy numbers of genes in fungi, aphids and other insects categorized by insect orders. The y-axis represents the average copy numbers per species (total copy numbers/total species). The total copy numbers are calculated based on the total protein sequences in each species. The species included are those analyzed in Fig. 2A. Each open circle represents a species.

**Supplementary fig. S3.**
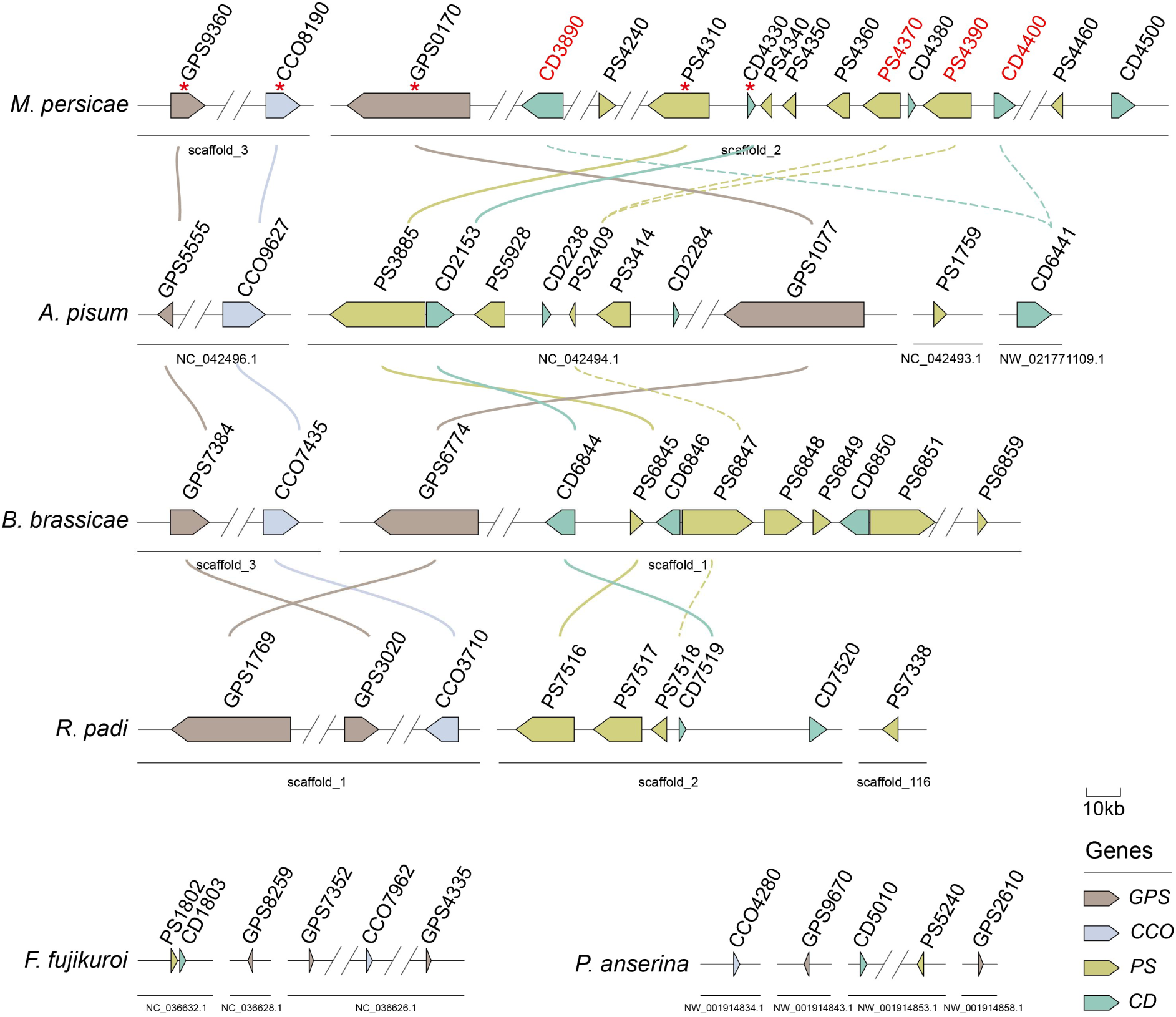
Genomic arrangement of carotenogenic genes in aphid species and fungi. This figure provides detailed information supplementing Fig. 3. Arrow directions indicate gene orientations, and double slashes represent manual breaks. Chromosomal positions are annotated below. *GPS*, *PS*, *CD*, and *CCO* genes are shown in different colors. Chromosome lengths are displayed proportionally, with a 10 kb scale bar. Red stars mark the collinear genes, while red gene IDs indicate duplicate genes in *M. persicae*. Collinear genes are connected by solid curves, and dashed lines link genes clustered with these duplicate genes.

**Supplementary fig. S4.**
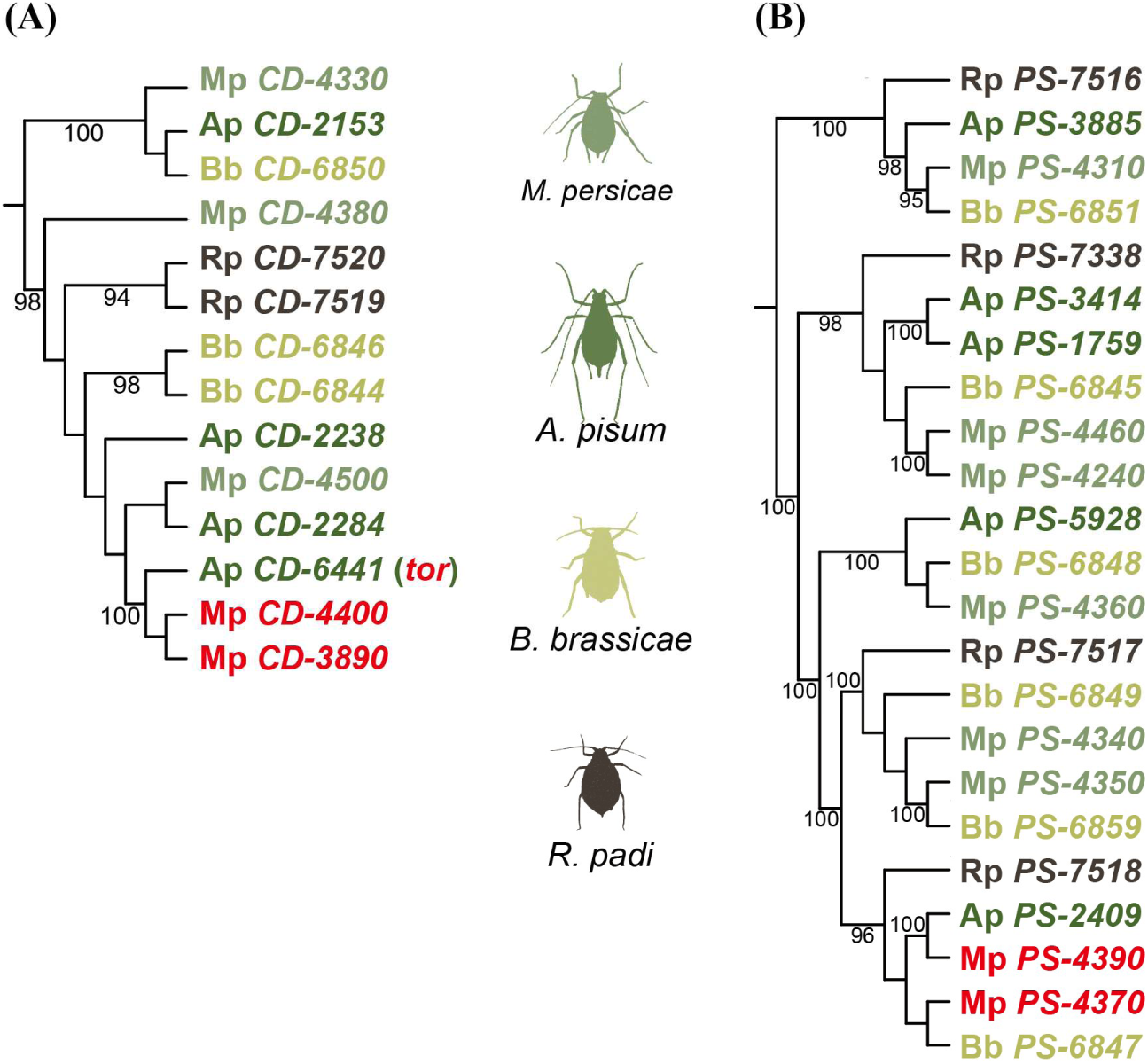
Phylogenetic trees illustrating duplicate genes and gene clusters among four aphid species. (A) tree of *CD* genes. (B) tree of *PS* genes. Gene IDs are marked with colors according to species. The gene Ap *CD-6441* is labeled as *tor*, a gene reported to associated with red body coloration in the pea aphid. Genes marked in red represent duplicates in *M. persicae*. Bootstrap values are shown on branches, with only values exceeding 90 displayed.

**Supplementary fig. S5.**
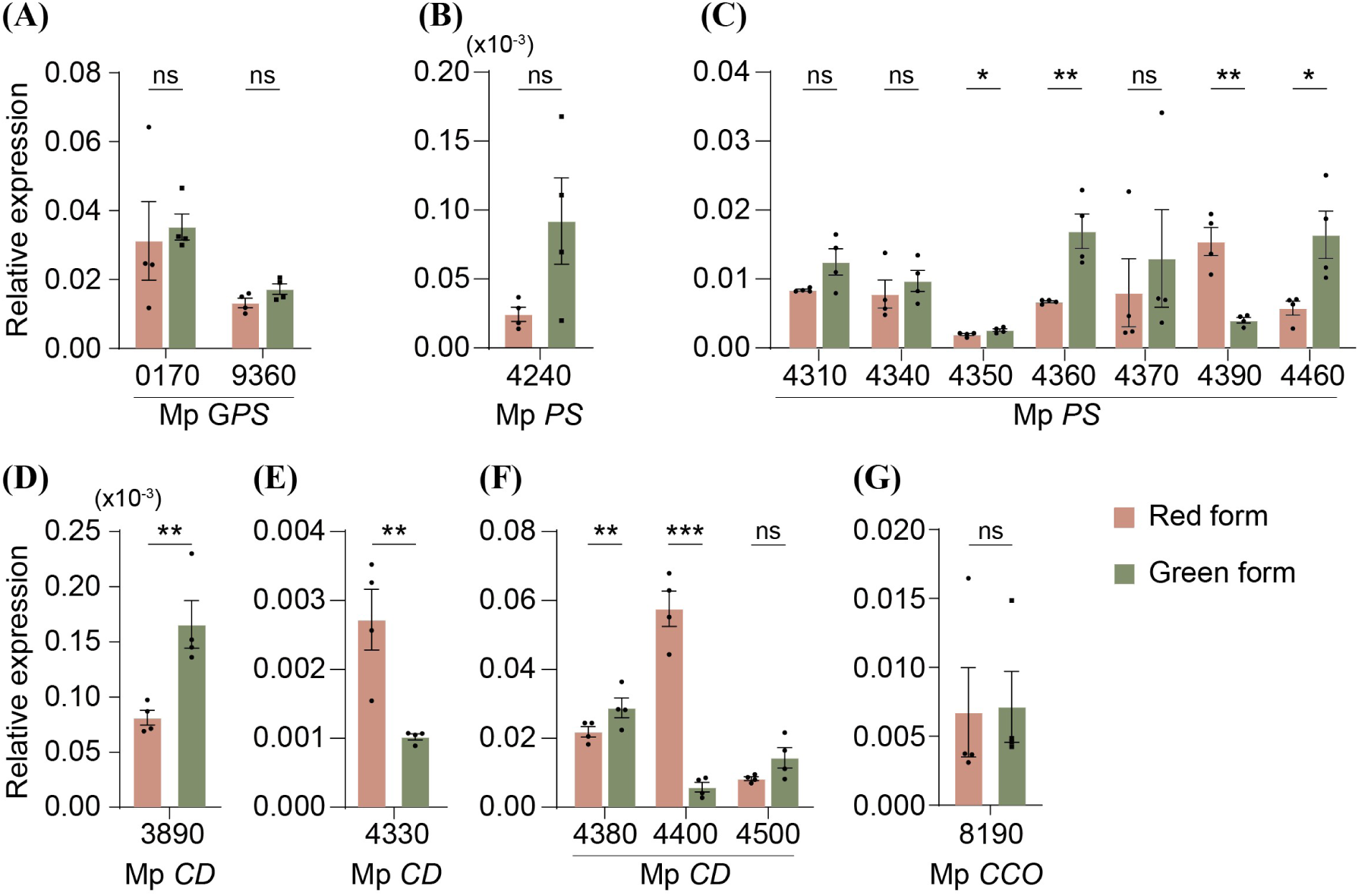
Gene expression comparison between the red form and green form aphids of the green-reddish clone. (A-G) Carotenogenic genes exhibit diverse expression patterns in the differently colored aphids. Solid circles represent replicates. (x10^-3^) denotes folded decimals for small Y-axis values. ns: not significant; *: *p*<0.05; **: *p*<0.01; ***: *p*<0.001. Unpaired two-tailed *t*-tests were used for all statistical analyses.

**Supplementary table S1.** The list of carotenogenic gene information from plant, fungi, and bacteria kingdoms.

**Supplementary table S2.** Genomic information of insects and the distribution of carotenogenic genes across insect species, along with the Pfam information for 288 insect species.

**Supplementary table S3.** Genome and Pfam information of plant, bacterial, and fungal species involved in this study.

**Supplementary table S4.** Primers used in this study.

**Supplementary table S5.** The detailed information of transcriptomic reads extracted from the NCBI SRA database.

